# Unifying turbulent dynamics framework distinguishes different brain states

**DOI:** 10.1101/2021.10.14.464380

**Authors:** Anira Escrichs, Yonatan Sanz Perl, Carme Uribe, Estela Camara, Basak Türker, Nadya Pyatigorskaya, Ane López-González, Carla Pallavicini, Rajanikant Panda, Jitka Annen, Olivia Grosseries, Steven Laureys, Lionel Naccache, Jacobo D. Sitt, Helmut Laufs, Enzo Tagliazucchi, Morten L. Kringelbach, Gustavo Deco

**Affiliations:** Computational Neuroscience Group, Center for Brain and Cognition, Department of Information and Communication Technologies, Universitat Pompeu Fabra, Barcelona, Catalonia, Spain; Universidad de San Andrés, Vito Dumas 284 (B1644BID), Buenos Aires, Argentina; Medical Psychology Unit, Department of Medicine, Institute of Neuroscience, University of Barcelona, Barcelona; Institute of Biomedical Research August Pi i Sunyer (IDIBAPS), Barcelona, Catalonia; Cognition and Brain Plasticity Unit, Bellvitge Biomedical Research Institute (IDIBELL), L’Hospitalet de Llobregat, Barcelona, Spain; Department of Cognition, Development and Educational Psychology, University of Barcelona, Barcelona, Spain; Institut du Cerveau et de la Moelle épinière, ICM, F-75013, Paris, France; Inserm U 1127, F-75013, Paris, France; CNRS UMR 7225, F-75013, Paris, France; Department of Neuroradiology, AP-HP, Hôpital Pitié-Salpêtrière, Sorbonne Université, 75006 Paris, France; Fundación para la Lucha contra las Enfermedades Neurológicas de la Infancia (FLENI), Buenos Aires, Argentina; Department of Physics, University of Buenos Aires. Intendente Güiraldes 2160 - Ciudad Universitaria - C1428EGA, Buenos Aires, Argentina; Coma Science Group, GIGA Consciousness, University of Liège, Liège, Belgium; Centre du Cerveau, University Hospital of Liège, Liège, Belgium; Department of Neurology, Christian Albrechts University, 24118 Kiel, Germany; Department of Neurology and Brain Imaging Center, Goethe University, 60528 Frankfurt am Main, Germany; Latin American Brain Health Institute (BrainLat), Universidad Adolfo Ibañez, Santiago, Chile; Department of Psychiatry, University of Oxford, Oxford, UK; Center for Music in the Brain, Department of Clinical Medicine, Aarhus University, DK; Centre for Eudaimonia and Human Flourishing, University of Oxford, Oxford OX1 2JD, United Kingdom; Institució Catalana de la Recerca i Estudis Avancats (ICREA), Barcelona, Catalonia, Spain; Department of Neuropsychology, Max Planck Institute for human Cognitive and Brain Sciences, Leipzig, Germany; School of Psychological Sciences, Monash University, Melbourne, Clayton VIC 3800, Australia

## Abstract

Recently, significant advances have been made by identifying the levels of synchronicity of the underlying dynamics of a given brain state. This research has demonstrated that unconscious dynamics tend to be more synchronous than those found in conscious states, which are more asynchronous. Here we go beyond this dichotomy to demonstrate that the different brain states are always underpinned by spatiotemporal chaos but with dissociable turbulent dynamics. We investigated human neuroimaging data from different brain states (resting state, meditation, deep sleep, and disorders of consciousness after coma) and were able to distinguish between them using complementary model-free and model-based measures of turbulent information transmission. Our model-free approach used recent advances describing a measure of information cascade across spatial scales using tools from turbulence theory. Complementarily, our model-based approach used exhaustive *in silico* perturbations of whole-brain models fitted to the empirical neuroimaging data, which allowed us to study the information encoding capabilities of the brain states. Overall, the current framework demonstrates that different levels of turbulent dynamics are fundamental for describing and differentiating between brain states.

## Introduction

Fundamentally different brain states such as sleep, wakefulness, or coma all emerge from the complex dynamics of self-organised brain activity. Nevertheless, an unanswered question in modern neuroscience is how best to characterise the underlying human brain states acquired with neuroimaging (Goldman et al., 2019; Kringelbach & Deco, 2020). Many challenges remain unsolved, and most importantly, there is a need to arrive at an agreed definition of brain states (Carhart-Harris et al., 2016; Deco et al., 2015; Gervasoni et al., 2004; Kringelbach & Deco, 2020; McCormick et al., 2020; Northoff, 2013; Tagliazucchi et al., 2016; G Tononi et al., 1994). The most important feature of such a definition would help to create a mechanistic framework for characterising brain states in terms of the underlying causal mechanisms and dynamical complexity. An elegant way of assessing dynamical complexity was proposed by Massimini and colleagues who investigated the perturbation-elicited changes in global brain activity during brain states, including wakefulness, sleep, anaesthesia, and post-coma states (Casali et al., 2013; Ferrarelli et al., 2010; Massimini et al., 2005). They have proposed the perturbational complexity index (PCI), which captures the significant differences in brain-wide spatiotemporal propagation of external stimulation, distinguishing between different brain states (Casali et al., 2013). Beyond basic neuroscience, a better definition and description of a brain state could offer novel avenues for translational therapeutic interventions to rebalance disrupted brain states in disease.

In a recent review, Goldman and colleagues (Goldman et al., 2019) showed that at both macroscopic and microscopic scales, unconscious brain states are dominated by synchronous activity (Brown et al., 2010; Fox, 2005; Henry, 2006; Sanchez-Vives & McCormick, 2000; Steriade et al., 1993), while conscious states are characterised by asynchronous dynamics (Boly et al., 2008; Raichle et al., 2001; Sanchez-Vives & McCormick, 2000). Equally, they propose that brain signals in unconscious and conscious states vary in their algorithmic complexity (Giulio Tononi & Edelman, 1998), entropy (Sitt et al., 2014), and dimensionality (El Boustani & Destexhe, 2010). The authors were inspired by the elegant mathematical framework of statistical physics, which provides the tools for uncovering structures of microscopic interactions underlying macroscopic properties. They propose that different brain states may emerge from the interactions between populations of neurons, similar to how different states of matter like solids and liquids emerge from interactions between populations of molecules. In other words, unconscious states are more like a solid-state, with high synchronicity and low complexity, while conscious states are more like liquids, with asynchronous activity and high complexity.

This dichotomy is very useful for capturing the fundamental difference between conscious and unconscious states, especially for the microstates, where for example, deep sleep is characterised by slow waves (Steriade, 2000). However, the transition between scales is more subtle and crucially depends on the complex percolation across the whole brain of the synchronous and asynchronous microstates, which gives rise to mixed complex dynamical states (Huber et al., 2004). The challenge remains to find a unifying dynamical approach, which can establish the balance between different levels of synchronicity and complexity needed to distinguish between brain states.

Here, we show that different brain states are always underpinned by spatiotemporal chaos, but the mixing across scales gives rise to dissociable turbulent dynamics. We investigate this using two complementary model-free and model-based frameworks. For the model-free approach, we profited from the immense advances in turbulence theory in physics (Frisch, 1995; Kolmogorov, 1941a, 1941b; Kuramoto, 1984). Recent advances have shown that the human brain operates in a turbulent regime (Deco, Kemp, et al., 2021; Deco & Kringelbach, 2020), which confers significant information processing advantages, including significant enhancing the functional role of the anatomical rare long-range connections (Deco, Perl, et al., 2021). This research has shown that brain information processing is favoured by turbulent dynamics, determined by the local synchronisation between brain areas. This level of local synchronisation can be linked to the rotational vortices found in fluid dynamics, and the size of these vortices defines the different scales of information processing.

For the model-based approach, we used whole-brain modelling based on the integration of anatomy and dynamics, which can be used to accurately fit and reproduce many aspects of empirical neuroimaging data (Breakspear, 2004; A. Ghosh et al., 2008; Honey et al., 2007; Jobst et al., 2017), and specifically to capture the brain turbulent dynamics (Deco & Kringelbach, 2020; Perl, Escrichs, et al., 2021). Crucially, this approach allows *in silico* exhaustive perturbation of the model that can be used to assess many aspects, including the susceptibility and information encoding capability. In other words, the model-free approach measures the naturally occurring information transmission flow, while the model-based approach allows us to measure the reactivity of the brain to external perturbations.

Here, we hypothesised that these two complementary measures will allow us to differentiate between different brain states. We found turbulent dynamics in all the different brain states but, crucially, using the model-free measure, we were able to characterise the different information transmission across spacetime scales in resting state, meditation, deep sleep and post-coma states. Furthermore, the model-based framework showed that different information encoding capabilities characterise different brain states. Thus, according to our hypothesis, the complementary methods are able to not only significantly distinguish between different brain states but also offer a unifying dynamical framework for mechanistically describing the underlying fundamental principles.

## Results

We used model-free and model-based frameworks to explore information transmission flow in whole-brain dynamics across different brain states. Specifically, we compared brain measures on three independent resting-state fMRI datasets. The meditation dataset comprised experienced Vipassana meditators (N=19) during both focused attention meditation (M) and resting state (R). The sleep dataset comprised healthy subjects during deep sleep, i.e., stage 3 (DS, N=13) and resting state (R, N=13) states. Finally, the disorders of consciousness (DOC) dataset were acquired in two independent research sites (Liège and Paris), comprised of healthy volunteers (R_CNT_: N=49) and DOC patients diagnosed in a minimally conscious state (R_MCS_: N=66) or an unresponsive wakefulness state (R_UWS_: N=39).

First, we applied the model-free approach to measuring information transmission flow across spacetime scales based on the recent finding demonstrating turbulence in human brain dynamics (**Fig. 1A**) (Deco & Kringelbach, 2020). This analysis was based on the local Kuramoto order parameter that describes the local level of synchronisation of a brain area, *n*, as a function of space, 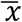, and time, *t*, at a given scale, *λ*. The scale of the local synchronisation is defined by the parameter *λ*, which determines the size of the spatial distances where the synchronisation is evaluated, where high values of *λ* stand for short distances, and vice versa (**Fig. 1B**). In particular, we computed for each dataset the amplitude turbulence (referred here as turbulence), information transfer, information cascade flow, and information cascade (**Fig. 1** and see more details in Methods).

**Fig. 1.**
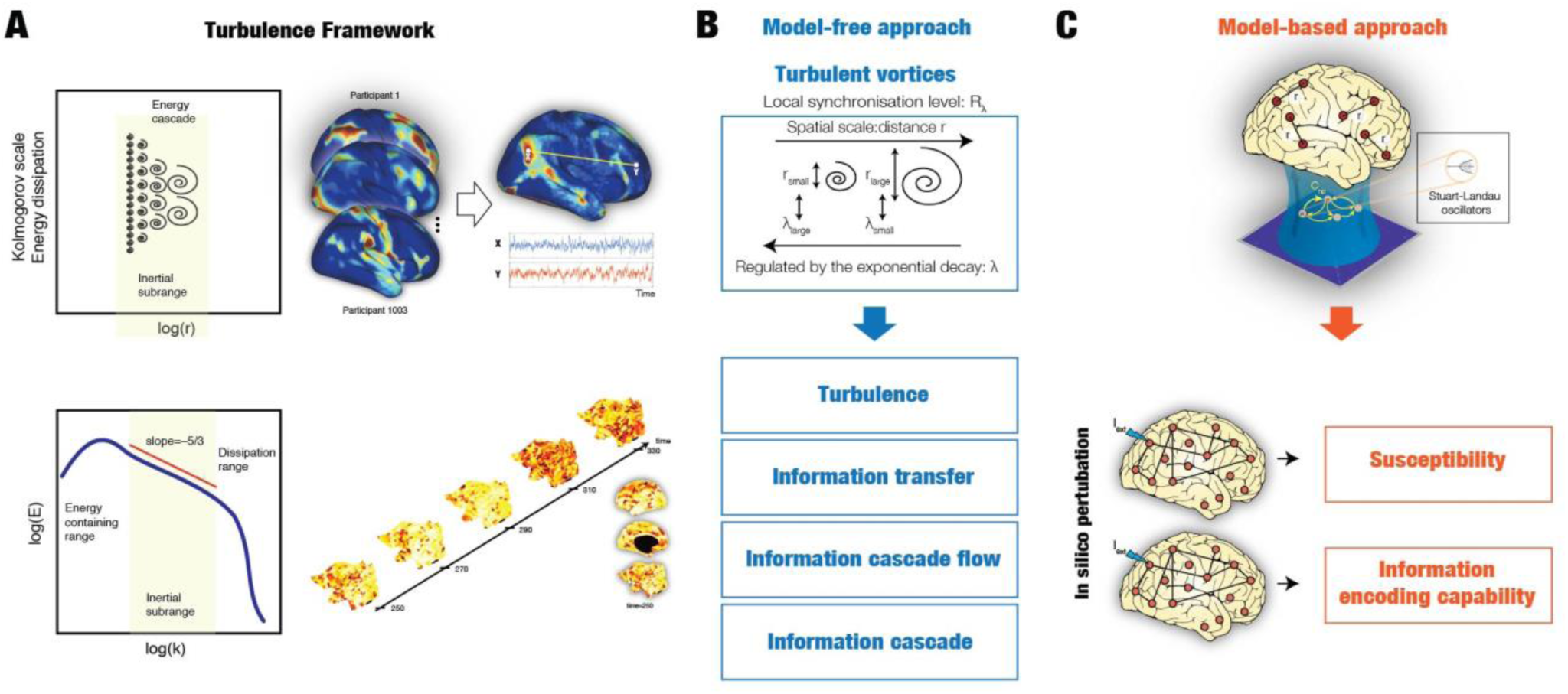
Overview of the empirical and model perturbative approach. **(A)** Turbulence in fluids is one the most common dynamical regime where the mixing motion governs (left panel). The energy cascade, i.e., how the energy travels across scale while dissipated (middle panel) and the statistical properties defined as power laws on the energy levels and structure functions (right panel) determine the turbulent behaviour of the fluid. (**B)** The analogy between brain activity and Turbulence can also be reflected in the similarity between the local level of synchronisation, determined by the local Kuramoto order parameter (R) at different scales (λ), and vortex with different spatial scales in fluid dynamics. The spatial scale (r) of the vortex is inversely related with the exponential decay of the local Kuramoto order parameter (λ). The turbulence regime also endows the brain with an efficient information cascade measured as the correlation of the local level of synchronisation across scales (information cascade flow). The average across scales of the information cascade flow is defined as the Information cascade. The information transfer quantified as the correlation of local synchronisation across space at different scales also characterises the brain’s information processing. (**C)** In the Hopf whole-brain model, the dynamics of each brain area are described through a Stuart Landau non-linear oscillator. The system of local oscillators is connected through the anatomical connectivity to simulate the global dynamics, capable of reproducing statistical observables from fMRI data. We used as structural connectivity the long-range connections (LR) from human diffusion MRI measurements on top of an exponential distance rule (EDR) to fit the empirical functional connectivity as a function of the Euclidean distance (following the relation between the Kolmogorov’s second-order structure-function and the traditional FC). Using whole-brain modelling allows obtaining measures that rise from the in silico perturbative approach. We simulated external stimuli and evaluated the model’s reaction for each brain state by quantifying the susceptibility and information capacity measures.

Second, we applied the model-based approach based on the sensitivity of these models to react to external *in silico* perturbations (**Fig. 1C** and Methods). For each brain state, we constructed a whole-brain dynamical model based on the normal form of a supercritical Hopf bifurcation coupled with the DTI structural connectivity and the exponential distance rule (EDR). Finally, to evaluate how each model fitted reacts to external stimuli, we applied *in silico* perturbations by quantifying the susceptibility and information encoding capability measures.

### Model-free framework

**Fig. 2** shows the results of the information transmission flow measures on the three datasets in terms of turbulence and information transfer. First, we explored the level of turbulence over different *λ* values, i.e., from 0.01 (∼100 mm) to 0.30 (∼3 mm), in steps of 0.03. This measure was defined as the standard deviation across time (*t*) and space (brain areas, *n*) of the local Kuramoto order parameter. Changes over time and space of this parameter are represented in the continuous snapshots rendered on a flatmap of the hemisphere for each condition in Fig. S1.

**Fig. 2.**
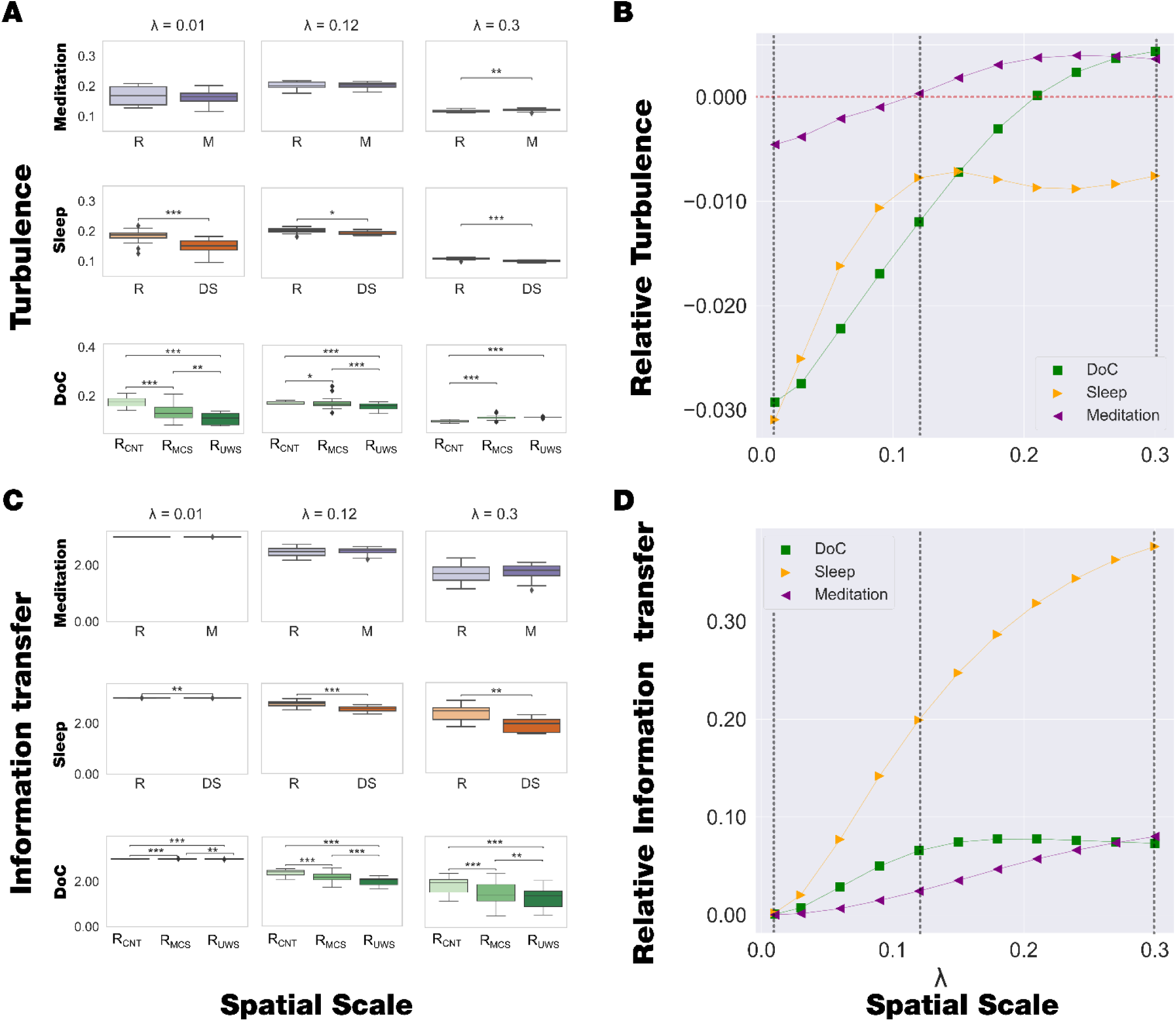
Model-free results: turbulence and information transfer. **(A)** We investigated the level of turbulence in different brain states, which measures the spatio-temporal variability of the local Kuramoto order parameter. We computed at different spatial scales from λ=0.01 (100 mm) to λ=0.03 (3mm) in steps of 0.03 and show the turbulence comparison between brain states within each dataset for λ=0.01, λ=0.12 and λ=0.3. The meditation state shows an increase in turbulence level compared to the resting state only on higher scales. By contrast, the deep sleep state shows lower turbulence than the resting state across all spatial scales, and turbulence decreases in the R_MCS_ and R_UWS_ states in lower lambda scales but increases in higher scales compared to healthy controls. **(B)** We computed a linear fit of the mean level of turbulence at each scale. For the liner fitting, we considered three brain states for DOC datasets (i.e., R_CNT_, R_MCS_, and R_UWS_) and two brain states for the sleep and meditation datasets (i.e., W, DS, and R, M, respectively). We display the obtained slopes as a function of the scale. The plot clearly shows that each dataset presented different behaviour. In particular, DOC showed negative slopes at lower scales and increased with the scales up to positive slopes. The sleep dataset presented negative slopes at lower scales, increased up to λ = 0.12, and a negative slope value kept almost constant. The meditation dataset also increased with scale but presented less variability than the other datasets. Dashed vertical lines indicate the scales displayed in panel A and the horizontal red dashed line highlights the zero slope. (**C)** We computed the information transfer, which measures how the information travels across space at different spatial scales. We display the comparison between brain states for λ=0.01, λ=0.12 and λ=0.3. The meditation state presents no significant differences on any scale compared to the resting state. In contrast, the information transfer significantly decreased for deep sleep and R_MCS_, R_UWS_ states compared to the resting state, across all scales. (**D)** We performed the same computation as panel B but considering the information transfer measure. In this case, the disorders of consciousness and sleep datasets presented a similar slope-scale relationship, whereas the meditation dataset presented less variability across scales. Dashed vertical lines indicate the scales displayed in panel C. P-values were assessed using the Wilcoxon rank-sum test and corrected for multiple comparisons, ∗ represents P<0.05, ∗∗ represents P<0.01 and ∗∗∗ represents P<0.001.

**Fig. 2A** shows the results of the turbulence level for each dataset. Specifically, the meditation state shows an increase in turbulence levels in higher spatial scales, i.e., short distances in the brain, compared to the resting state. On the other hand, the deep sleep state shows lower turbulence levels than the resting state across all the spatial scales. Finally, the turbulence levels decrease for DOC patients (R_MCS_ and R_UWS_) compared to healthy controls during resting state in lower lambda values, i.e., long distances, but increases in higher lambda scales; differentiating, also, between the R_UWS_ and R_MCS_ groups. To summarise the behaviour of the time and space information transmission measures at different scales, we quantified the turbulence changes at each *λ* across brain states. We computed a linear fit to the mean turbulence of brain states at each *λ* and obtained the slopes of the corresponding lines, which stands for turbulence across brain states at a specific scale. **Fig. 2B** shows the relationship between these slopes and scales for each dataset. The meditation dataset presents similar behaviour but is less sensitive to this measure, i.e., lower variability of the slope values across scales. By contrast, the sleep dataset shows a monotonical increase of slope values from negative values at low scales up to *λ* = 0.12, where it remains almost constant for higher scales. Finally, DOC states present the same behaviour: the slopes monotonically increase from negative values at low *λ* scales towards positive values at high *λ*. It is noticeable that with this quantification it is possible to differentiate between datasets that involve a reduction of consciousness, i.e., despite that sleep and DOC patients present a reduction of the information processing in many scales, the behaviour across scales captures differences between sleep and DOC states.

**Fig. 2C** displays the results of the information transfer, which indicates how the information travels across space at a given spatial scale, *λ*. The information transfer in the meditation state shows no significant differences across any scale compared to the resting state. By contrast, this measure significantly decreases for deep sleep and DOC states across all *λ* scales compared to the resting state, and interestingly, differentiating the MCS and UWS groups across all scales.

To summarise the behaviour of information transfer at different scales, we quantified the changes at each *λ* across brain states. Conversely, the evolution of the slopes across scales for the information transfer presents the same behaviour across all datasets (**Fig. 2D**).

**Fig. 3A** shows the information cascade flow across scales, which indicates how the information travels from a given scale, *λ*, to a lower scale, *λ* − Δ*λ*, in consecutive time steps, *t* and *t* + Δ*t*. This measure presents no significant differences in the meditation state compared to the resting state, whereas for deep sleep and DOC, the information cascade flow decreases across all scales compared to the resting state.

**Fig. 3.**
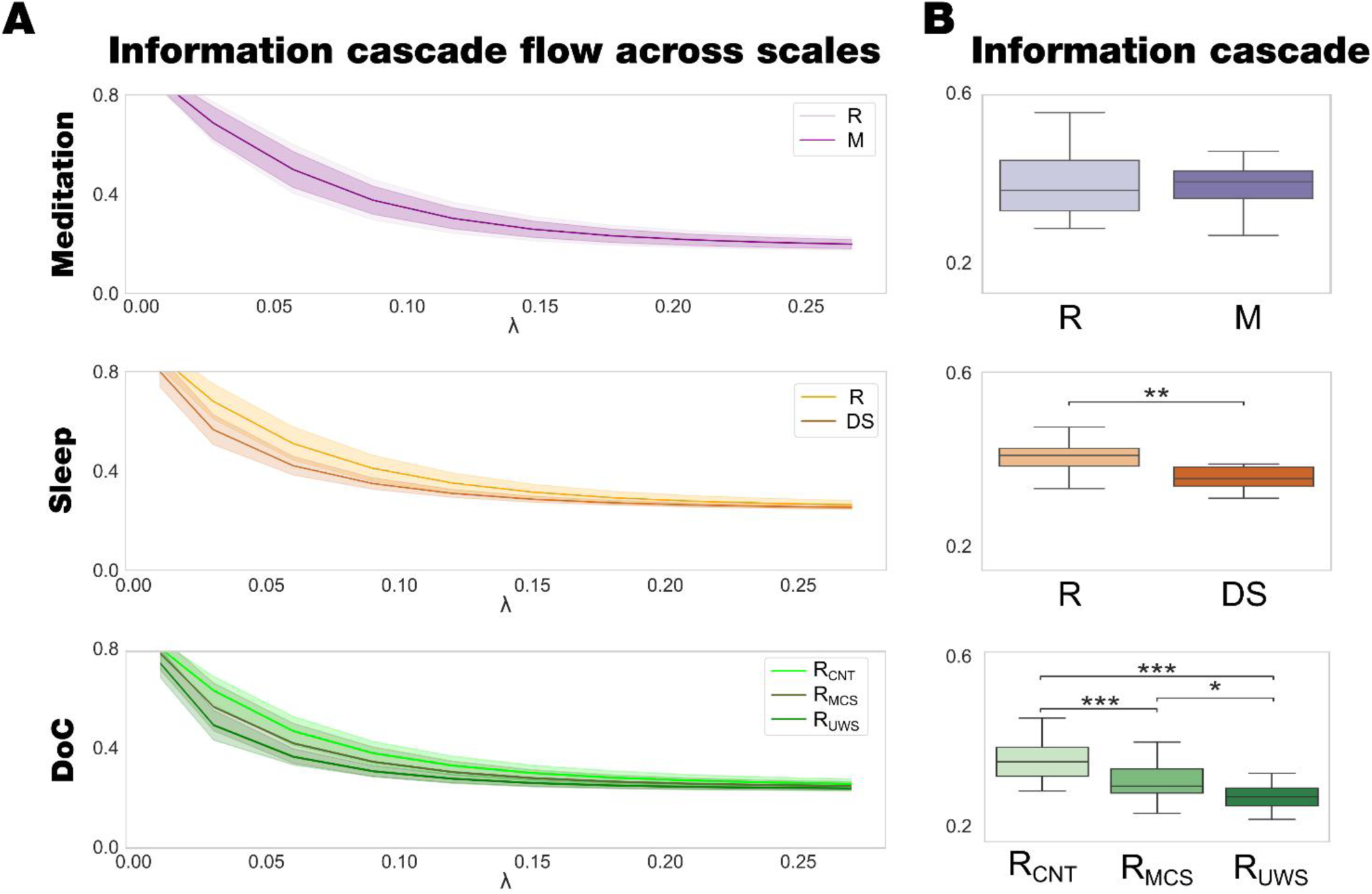
Model-free results: information cascade flow and information cascade. **(A)** The information cascade flow across scales is the predictability given by the level of synchronisation at a specific scale (λ) from the previous scale λ−Δλ (where Δλ = 0.03 is the discretisation of scale). The meditation state presents no differences across the scales compared to the resting state, the information cascade flow significantly decreases for DS and R_MCS_, R_UWS_ states compared to the resting across all scales. (**B)** The information cascade, defined as the average information cascade flow, differentiates R_MCS_, R_UWS_, and DS states from the resting state, while the meditation state presents no differences. P-values were assessed using the Wilcoxon rank-sum test and corrected for multiple comparisons, ∗ represents P<0.05, ∗∗ represents P<0.01 and ∗∗∗ represents P<0.001.

**Fig. 3B** shows the information cascade. This measure is obtained by averaging the information cascade flow across all λ scales and summarises the whole behaviour of the information transmission across scales. The information cascade in the meditation state presents no significant differences compared to the resting state. In contrast, the deep sleep and DOC states present less information transfer across the scales than the resting state, moreover, the information cascade clearly differentiate between R_CNT_ and R_UWS_ states.

### Results of node-level turbulence in different brain states

We computed the node-level turbulence as the standard deviation over time of the local Kuramoto order parameter for each brain state in each dataset. This measure indicates how changes the level of local synchronisation across time. **Fig. 4** shows the results of the node-level turbulence in different brain states (resting state, meditation, deep sleep, and DOC). We quantified this difference by computing the Kolmogorov-Smirnov distance (KSD) between the distributions of node-level turbulence, where larger values mean more different distributions (see Methods).

**Fig. 4.**
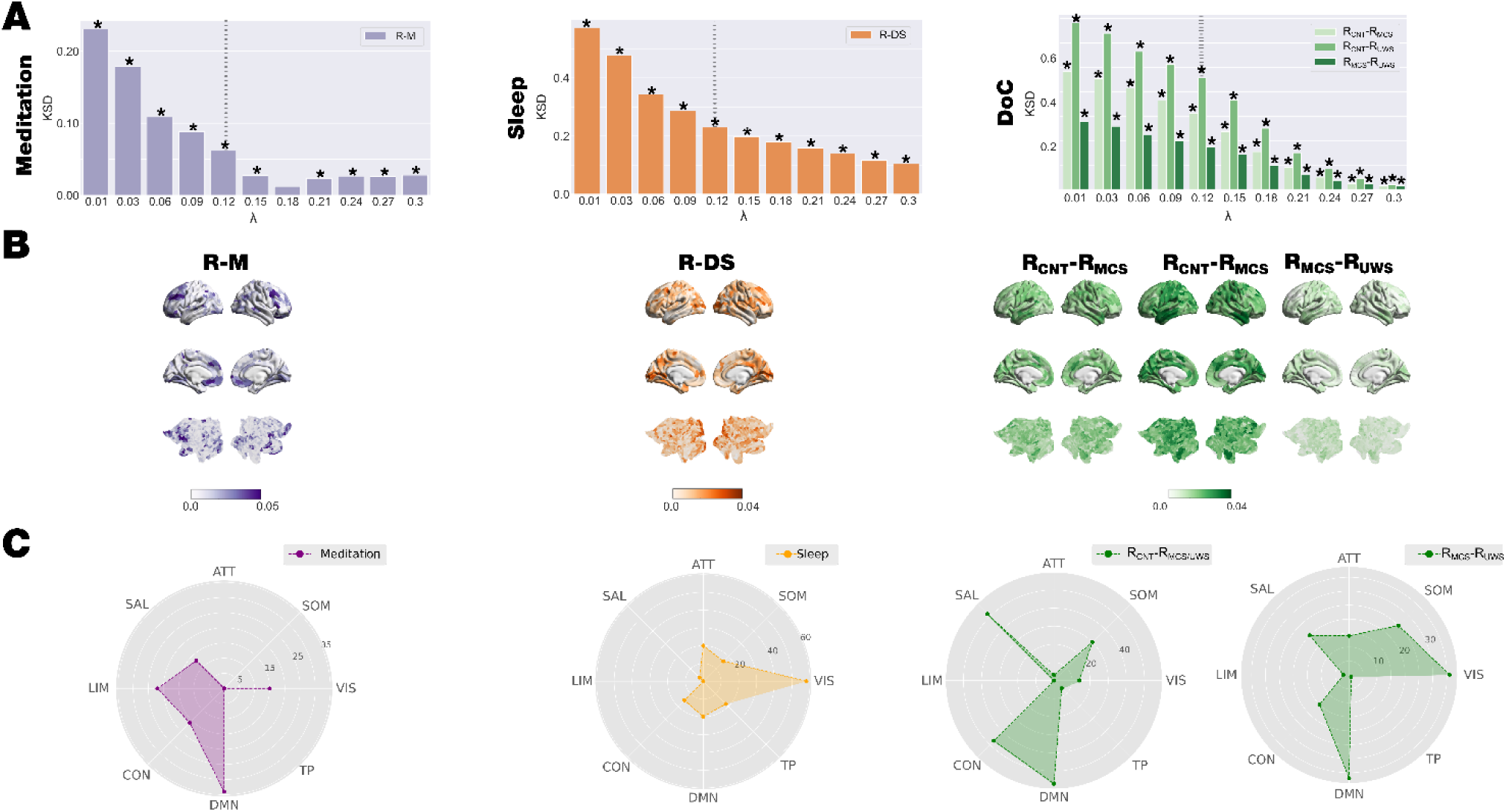
Model-free results: Node-level turbulence. We computed the node-level turbulence as the standard deviation across time of the local Kuramoto order parameter. **(A)** We performed the KSD between distributions of the node-level of turbulence of each brain state within each dataset for each scale. The KSD for all datasets monotonically decreases, whereas the value of λ increases for all comparisons. **(B)** Render brains represent the absolute difference of the node-level turbulence between each brain state for scale λ = 0.12, indicated with vertical dashed lines in panel A. We selected the top 15% quantile of absolute differences between conditions, identified the resting state networks to which they belong and quantified the number of nodes per network. **(C)** Radar plots represent the number of nodes on the top 15% quantile of the absolute difference by each comparison and resting-state network (CON: control; DMN: default mode; TP: temporal-parietal; VIS; visual; SOM: somatomotor; ATT: attentional; SAL: salience; LIM: limbic). The networks showing the highest differences between resting and meditation states were the limbic and default-mode networks. The comparison between deep sleep and resting state shows that nodes of the visual- and default-mode-networks present the highest difference. Finally, the comparison between R_CNT_ and DOC patients (R_MCS_ and R_UWS_) shows that the somatomotor-, salience-, control-, and default-mode-networks present the highest differences, whereas, specifically in the comparison between R_MCS_ and R_UWS_ nodes associated with the somatomotor- and control-networks present the highest differences. P-values were assessed using the Kolmogorov– Smirnov test and corrected for multiple comparisons, ∗ represents P<0.001.

**Fig. 4A** displays the KSD between brain states across scales. The KSD for all datasets monotonically decreases, whereas the value of λ increases. In other words, the KSD is maximal for lower values of λ, i.e., long distances in the brain. In particular, for DOC states, the higher KSD is found between R_CNT_ and R_UWS_ states.

**Fig. 4B** shows the absolute difference between the node-level turbulence between brain states in each dataset at *λ* = 0.12 rendered onto the brain cortex. First, we selected the nodes for each comparison of the top 15% quantile. Then, we identified the resting state networks to which they mainly belong and quantified the number of nodes per network (**Fig. 4C)**. We found that differences between meditation and resting state are mainly in the limbic and default-mode networks. In contrast, the highest differences between deep sleep and resting state are observed in the nodes of the visual- and default-mode-networks. Finally, the highest differences in local synchronisation are found between controls during resting state and DOC patients (R_MCS_ and R_UWS_) in the somatomotor-, salience-, control-, and default-mode-networks. Conversely, the highest differences between R_MCS_ and R_UWS_ are observed in nodes associated with visual-, somatomotor- and default mode-networks.

### Model-based framework

For each brain state, we built a Hopf whole-brain model of coupled dynamical oscillators in an anatomical brain architecture coupling the exponential distance rule (EDR) and the DTI matrix fitted to the empirical functional data (see more details in Methods).

**Fig. 5A** shows the functional connectivity fitting as a function of the global coupling parameter, *G*, for each dataset. For each *G*, we repeated 100 simulations for each brain state with the same TR and time duration as the empirical data. Then, we computed the fitting of the functional connectivity as the Euclidean distance between the empirical and simulated functional connectivity (FC). The optimal working point of each model is determined as the minimum of the fitting level (vertical lines in **Fig. 5A**). We used the respective minima of each condition as the basis of the following perturbative *in silico* investigations. The *G* values obtained for meditation, deep sleep, and DOC are lower than those obtained for the resting state. This result can be interpreted as reducing the coupling between areas to represent the global brain dynamics.

**Fig. 5.**
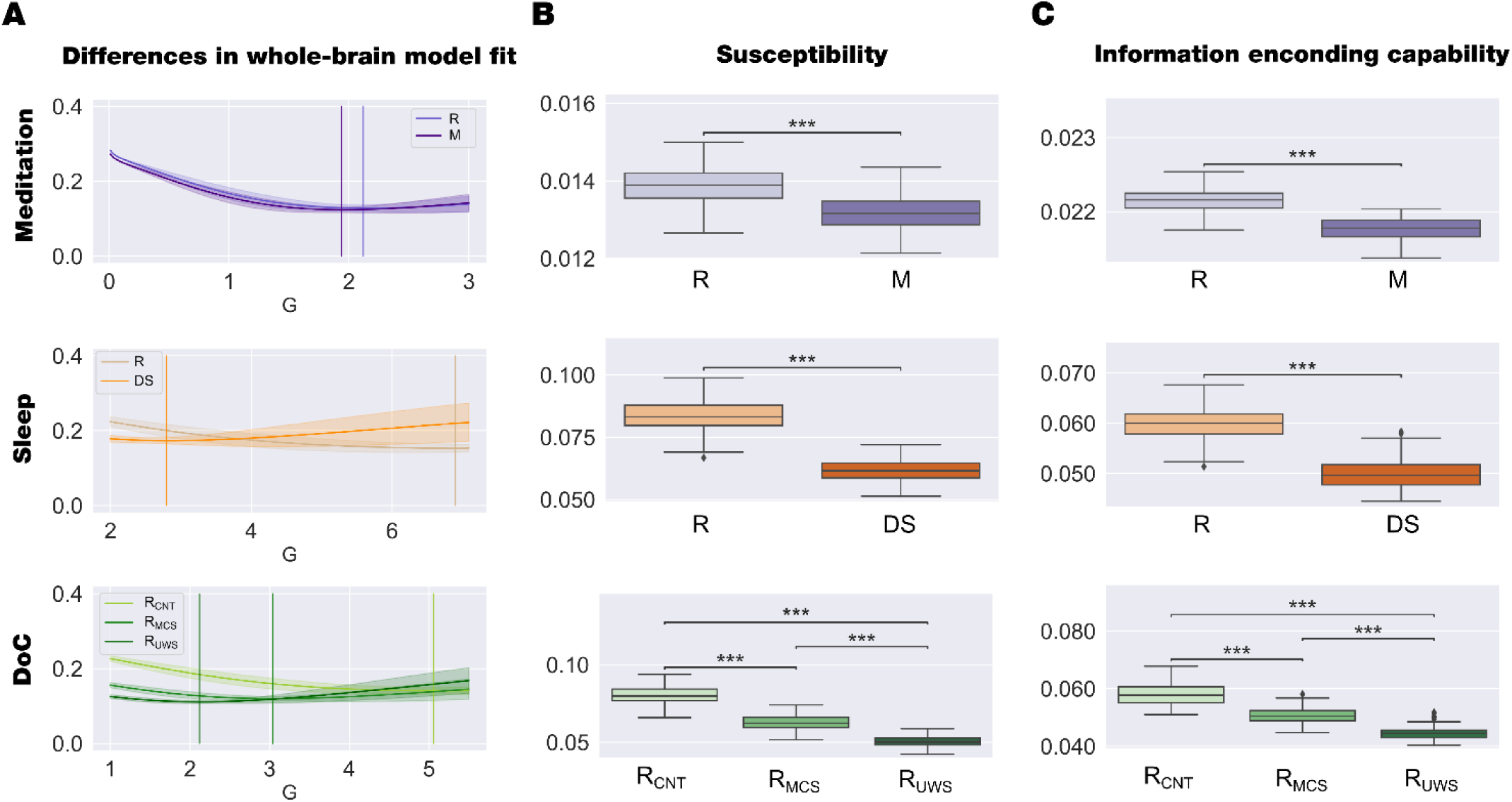
Model-based perturbative approach. (**A)** We show the evolution of the error of the whole-brain model FC fitting to the empirical fMRI data as a function of the global coupling strength, G. The error of the FC fitting was given by the square root of the difference between the simulated and empirical FC matrix. The optimal working point of the model was defined as the minimum value of the FC fitting, i.e., where the model shows maximal similarity to the empirical fMRI data. (**B)** We show the results of the susceptibility measure, which estimates how these models react to external artificial perturbations. In all datasets, the resting state was the most susceptible to be perturbed. (**C)** We show the information encoding capability of the whole-brain models, which captures how different external stimulations are encoded in the dynamics. Similar to the susceptibility measure, the resting state was more susceptible to react to the perturbations. Susceptibility and information capacity measures differentiated each brain state and between R_MCS_ and R_UWS_ groups. These results show that each brain state encodes the whole-brain dynamics with a particular complexity. P-values were assessed using the Wilcoxon rank-sum test and corrected for multiple comparisons; ∗∗∗ represents P<0.001.

**Fig. 5B-C** displays the results of perturbing each whole-brain model using *in silico* stimulations. To study how the system reacts to external perturbations, we perturbed each model at its optimal working point and computed the model-based measures. Specifically, we estimated the susceptibility by measuring the perturbed and non-perturbed modulus of the local Kuramoto order parameter 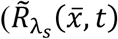 and 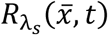, respectively). The perturbation consisted of randomly changing the bifurcation parameter of each brain area, *a*_*n*_, within the range [-0.02:0] (**Fig. 1C**). Finally, we calculated the susceptibility as the difference between the perturbed and non-perturbed cases averaged across time, trials, and space. **Fig. 5B** compares the distribution of susceptibility computed in 100 repetitions of the *in silico* perturbative approach for each condition within each dataset. Similarly, we computed the information encoding capability (as an extension of the susceptibility) to study how external perturbations are encoded in brain dynamics. This measure was defined as the standard deviation across trials of the difference between the perturbed and unperturbed mean of the modulus of the local Kuramoto order parameter across time, averaged across space. **Fig. 5C** shows the results obtained for each condition within each data set. Both measures show that the capability to react under external perturbations decreases for meditation, deep sleep, and DOC compared to the resting state.

## Discussion

Here, we have proposed and tested a unifying turbulent dynamics framework for defining and distinguishing between different brain states. We used model-free and model-based approaches to demonstrate that different levels of turbulent dynamics and information encoding can differentiate between brain. These approaches are based on the recent finding demonstrating turbulence-like dynamics in the healthy human brain (Deco, Kemp, et al., 2021; Deco, Perl, et al., 2021; Deco & Kringelbach, 2020). Crucially, our model-free approach was able to show the role of information cascade across spatial scales as a distinguishing feature between brain states. Specifically, we investigated information transmission across different spatial scales by analysing neuroimaging fMRI data during resting state, meditation, deep sleep, MCS, and UWS. Our results showed that these brain states exhibit significant differences in information cascade across different scales at both the spatial and temporal domains. Equally, our model-based approach using exhaustive *in silico* perturbations of whole-brain models demonstrated the role of susceptibility and information encoding capabilities between different brain states. The results illustrated that by inducing a shift in the intrinsic local dynamics of brain areas, the brain responds to the external perturbations less sensitively as the conscious awareness diminishes. This unifying framework captures the percolation between scales across the whole brain of the different levels of synchronicity and complexity found in microstates associated with different brain states. At the macroscale, the result of this percolation and mixing across scales is always spatiotemporal chaos. However, there is a clear structure in this chaos, which is well described by different levels of turbulent dynamics. Overall, the proposed unifying framework reconciles the balance between different levels of synchronicity and complexity in the turbulent framework of spatiotemporal chaos for describing and differentiating between brain states. Importantly, both model-free and model-based measures successfully differentiate the minimally conscious state (R_MCS_) and unresponsive wakefulness syndrome (R_UWS_) groups.

### Brain states are characterised by different turbulence dynamics across scales

Previous research has demonstrated that the turbulent regime in brain dynamics enhances information processing of long-range connections in the resting state of healthy awake participants (Deco, Perl, et al., 2021). However, we found that the level of turbulence is a sensitive measure to distinguish between different brain states. In particular, we found that the evolution of turbulence and information cascade at different spatial scales is a fundamental feature of the different brain states. In particular (as shown in **Fig. 2A** and **2B**), when compared to healthy controls, the DOC groups showed lower levels of turbulence at higher spatial scales (i.e., lower λ and larger distances) but higher levels of turbulence at lower spatial scales. Crucially, long-range information transmission has been shown to be associated with cognition in healthy participants (Deco, Perl, et al., 2021) and its dramatic reduction found in UWS and MCS patients (see relevant boxplot for λ=0.01 in **Fig. 2A**) suggests that this could be a defining feature of the reduction of consciousness. This result is consistent with EEG evidence showing that noncommunicative patients have lower global information sharing (King et al., 2013), and decreased brain complexity (Bodart et al., 2017; Casarotto et al., 2016). Our findings are also aligned with the global neuronal workspace theory that postulates that the long-distance connexions globally broadcast the information for different processor brain-wide and this lack of spatially bounded information processing is associated with conscious (Dehaene & Changeux, 2011).

In healthy participants, the unconscious state of deep sleep was characterised by lower turbulence across all spatial scales, demonstrating a reduction in information processing over both short and long distances (Tagliazucchi et al., 2013; Tagliazucchi & van Someren, 2017). In contrast, in highly trained meditators, the meditation state presented higher turbulence only at lower spatial scales (large λ and short distances), suggesting that meditation is a state showing an alteration rather than a reduction of consciousness.

Overall, we have demonstrated that each brain state exhibits different turbulent dynamic patterns across spatial scales, allowing us to characterise the brain states from their underlying information processing fluctuations. Interestingly, it allowed us to differentiate between deep sleep and DOC states, thus unveiling specific and unique features of turbulent dynamics underlying low-level states of consciousness.

### Brain states are characterised by different information transmission across scales

Given that different turbulent regimes can describe different brain states, we were interested in determining how this impacts the underlying information processing across scales. To this end, we quantified three different measures of information transfer, information cascade flow and information cascade for each brain state.

As shown in **Fig. 2C** and **2D**, the differences in information transfer increase with the spatial scale between normal resting state and the level of awareness in meditation, deep sleep and DOC. This result clearly demonstrates that the measure of information transfer indexes conscious awareness. Interestingly, while this measure increases with the distance between resting and meditation, this difference is not statistically significant. This suggests that meditation is more like the resting state but with important significant differences revealed by the other information transmission measures.

The information cascade flow monotonically decreases with lower distance (the increase in spatial scale λ) for all brain states (shown in **Fig. 3A**). This measure also discriminates *between* conditions within each dataset, i.e., showing lower values for DOC patients than control participants and in the deep sleep stage compared to the resting state in the same participants.

The information cascade (i.e., the average of the information flow across scales) is lower in low-levels states of awareness (deep sleep, R_MCS_ and R_UWS_) than in normal resting state (shown in **Fig. 3B**).

Overall, this demonstrates that the information transmission is altered with conscious awareness and that this is captured with the global information processing measures of information transmission, information cascade flow and information cascade.

### Fine-grained node-level turbulence differences between brain states

In contrast, it is also possible to investigate the local changes in turbulence between different brain states to identify local regions potentially involved in controlling the turbulent dynamics. To this end, we defined a local measure of the regional level of variability of the local node-level of turbulence. We found that this measure is also able to differentiate between different brain states at different spatial scales. In particular, our analysis of the node-level local synchronisation shows that higher *λ* values, i.e., shorter distances in the brain, are less sensitive in discriminating between brain states (**Fig. 4A**).

In addition, the node-level of description allows us to discriminate the brain areas involved in the whole-brain dynamic changes between brain states. As shown in **Fig. 4B** (at *λ*=0.12), we found that brain regions belonging to the somatomotor, salience, control, and default-mode networks present the most critical differences in DOC states, with a more substantial decrease in the R_UWS_ than in the R_MCS_ condition, corresponding to lower levels of conscious awareness. Specifically, we found the highest difference between R_UWS_ and R_MCS_ in brain regions belonging to default mode-, visual- and somatomotor-networks, which is consistent with previous studies in DOC patients (Bodien et al., 2017; Athena Demertzi et al., 2015; Fernández-Espejo et al., 2012; Qin et al., 2015). We also found that changes in regions in visual- and default-mode-networks indexed differences between deep sleep and wakeful resting, consistent with other studies of the human wake-sleep cycle (Horovitz et al., 2009; Ipiña et al., 2020; Stevner et al., 2019). In contrast, comparing meditation with resting state in expert meditators revealed regions in limbic- and default-mode-networks, similar to other findings in meditation (Filippi et al., 2021; Hasenkamp et al., 2012; Hasenkamp & Barsalou, 2012; Tang et al., 2015; Taylor et al., 2013).

### Revealing susceptibility and information encoding capability using whole-brain modelling

Given the exciting results of directly perturbing the brain revealed by the pioneering studies of Massimini and colleagues (Casali et al., 2013; Ferrarelli et al., 2010; Massimini et al., 2005), we also wanted to explore the causal mechanistic underpinnings of the differences between brain states and ensuing reactivity to external perturbations. To this end, we modelled the empirical fMRI data using Hopf whole-brain models (Deco & Kringelbach, 2020; Anandamohan Ghosh et al., 2008; Ipiña et al., 2020; Jobst et al., 2017). We found that the optimal working point of the whole-brain models for all brain states shifted to a lower global coupling factor compared to the resting state (see **Fig. 5A**). The global coupling parameter, *G*, represents the conductivity of the fibre densities among brain regions given by the underlying structural connectivity, which is assumed, for simplicity, to be equal across the brain (Deco, Kringelbach, et al., 2017; Deco, Tagliazucchi, et al., 2017). A higher coupling, *G*, allows the propagation of information among brain areas indirectly connected, enhancing the transmission of information across the whole network and vice versa (López-González et al., 2021). Overall, this drastic shift toward a lower coupling indicates sub-critical behaviour suggestive of a change in the dynamical complexity underlying the brain state (Deco, Tagliazucchi, et al., 2017). These results are in line with previous studies showing a reduced spatiotemporal dynamic across the whole-brain functional network in DOC patients (A. Demertzi et al., 2019; Hannawi et al., 2015; López-González et al., 2021; Schiff et al., 2014), deep sleep (Horovitz et al., 2009; Ipiña et al., 2020; Jobst et al., 2017; Schartner et al., 2017), and meditation (Escrichs et al., 2019; Toutain et al., 2021) compared to normal resting state.

Crucially, we perturbed each brain model at its optimal working point to investigate the induced whole-brain dynamics changes caused by the external *in silico* perturbations. Specifically, our external manipulation consisted of a shift towards the bifurcation point of the intrinsic local dynamics of brain areas. We found that the resting state shows significantly higher susceptibility and information encoding capability than in the pairwise comparison in each dataset, i.e., meditation, deep sleep and DOC. The similar behaviour of both measures (susceptibility and information encoding capability) can be related to the specific features of our perturbative approach. Differences *in silico* protocols can be assessed to study how different brain states react to external perturbations such as shifting the local dynamics in the opposite direction, node by node perturbation (Deco et al., 2019), non-sustained perturbations (Deco et al., 2018; Perl, Escrichs, et al., 2021) or perturbing with external periodic force (Perl, Escrichs, et al., 2021; Perl, Pallavicini, et al., 2021). Notably, the perturbative approach allows for the exploration of brain responses elicited by *in silico* protocols which are not limited by ethical constraints of *in vivo* stimulations (Clausen, 2010; Kringelbach et al., 2007). Furthermore, the differential sensitivity of each brain state of external perturbations can serve as a specific biomarker that reveals features of their dynamical complexity.

Overall, we have presented a unifying framework that can account for the differences between brain states. The key idea is that the complex dynamics of a brain state result from the percolation across scales of previously demonstrated differences in synchronicity and complexity at the microscale. These dynamics are always spatiotemporal chaos but with differentiable turbulent dynamics, which our dual model-free and model-based approach can reveal. The main finding is that turbulent dynamics across different spatial scales can distinguish between brain states. Furthermore, these differences are also found as differences in susceptibility and information encoding capability as a result of the reactivity of different external perturbations on the underlying brain state. Given the sensitivity and specificity of the results, long-term, these might help identify potential targets for patients to rebalance and regain consciousness.

## Methods

### Participants

#### Meditation

A total of 20 experienced meditators with more than 1000 hours of meditation experience were selected from a dataset previously described in Escrichs et al. (2019). Meditators were recruited from Vipassana communities of Barcelona, Catalonia, Spain (7 females, mean ± SD, 39.8±10.29 years, 9,526.9±8,619.8 meditation experience). Participants were asked to practice focused attention on breathing (i.e., anapanasati in language Pali). In this meditation technique, meditators focus their attention on natural breathing, and when they realize that the mind is wandering, they must refocus their attention back to natural breathing. All participants reported no history of past neurological disorder and gave written informed consent. The study was approved by the Ethics Committee of the Bellvitge University Hospital according to the Helsinki Declaration on ethical research.

#### Sleep

A total of 63 healthy subjects (36 females, mean ± SD, 23±43.3 years) were selected from a dataset previously described in Tagliazucchi and Laufs (Tagliazucchi & Laufs, 2014). On the day of the study, participants reported a wake-up time between 5:00 AM and 11:00 AM and a sleep onset time between 10:00 PM and 2:00 AM for the night before the experiment. Within half an hour of 7 PM, participants entered the scanner and were asked to relax, close their eyes, and not fight the sleep onset. Their resting state activity was measured for 52 minutes with a simultaneous combination of EEG and fMRI. According to the rules of the American Academy of Sleep Medicine (Berry et al., 2015), the scalp potentials measured with EEG determine the classification of sleep into four stages (resting state, N1, N2 and N3 sleep). We selected 13 subjects who reached the deep sleep stage (DS, i.e., N3) and contiguous time series of at least 198 volumes. The local ethics committee approves the experimental protocol (Goethe-Universität Frankfurt, Germany, protocol number: 305/07), and written informed consent was asked to all participants before the experiment. The study was conducted according to the Helsinki Declaration on ethical research.

#### Disorders of consciousness: Paris

A total of 77 patients who were hospitalized in Paris Pitié-Salpêtrière, suffering from brain injuries, were included in this study. Clinical assessment and trained clinicians carried out the clinical assessment and Coma Recovery Scale-Revised (CRS-R) scoring to determine their state of consciousness. Patients were diagnosed with UWS if they showed arousal (opening their eyes) without any signs of awareness (never exhibiting non-reflex voluntary movements). On the other hand, patients were in a MCS if they exhibited some behaviours that could be indicative of awareness, such as visual pursuit, orientation to pain, or reproducible command following. We excluded subjects with T1 acquisition errors (n=5), with high levels of motion detected (n=7), registration errors (n=4), and large focal brain lesions (n=4). We thus included 33 patients in MCS (11 females, mean age ± SD, 47.25± 20.76 years), and 24 in UWS (10 females, mean age ± SD, 39.25± 16.30 years) and 13 healthy controls (7 females, mean age ± SD, 42.54± 13.64 years). This research was approved by the local ethics committee Comité de Protection des Personnes Ile de France 1 (Paris, France) under the code ‘Recherche en soins courants’ (NEURODOC protocol, n° 2013-A01385-40). The patient’s family gave their informed consent for the participation of their relative, and all investigations were conducted according to the Declaration of Helsinki and the French regulations.

#### Disorders of consciousness: Liège

A total of 35 healthy controls (14 females, mean age ± SD, 40 ± 14 years) and 48 patients with disorders of consciousness (DOC) were included in the study based on a dataset previously described in López-González et al (López-González et al., 2021). The diagnosis was made after at least 5 CRS-R by trained clinicians. The highest diagnosis of the level of consciousness was taken as the final diagnosis, which was also confirmed with Positron Emission Tomography (PET) (i.e., patients in MCS presented a relatively preserved metabolism in the fronto-parietal network while patients with UWS had a bilateral hypometabolism in this network). We thus included 33 patients in MCS (9 females, mean age ± SD, 45 ± 16 years), and 15 in UWS (6 females, mean age ± SD, 47 ± 16 years). The Ethics Committee of the Faculty of Medicine of the University of Liege approved the study protocol. The study was conducted according to the Helsinki Declaration on ethical research. Written informed consent was obtained from controls and the patients’ legal surrogates.

### MRI Data Acquisition

#### Meditation

MRI images were acquired on a 3T Siemens Trio scanner (Siemens, Erlangen, Germany) using a 32-channel receiver coil. The high-resolution T1-weighted images were acquired with 208 contiguous sagittal slices; TR/TE= 1970 ms/ 2.34 ms; inversion time (IT) = 1050 ms; flip angle = 9°; FOV = 256 mm; and isotropic voxel size 1 mm. Resting-state and meditation fMRI images were performed by a single shot gradient-echo EPI sequence with a total of 450 volumes (15 min); TR/TE = 2000 ms/29 ms; FOV= 240 mm; in-plane resolution 3 mm; 32 transversal slices with thickness = 4 mm; flip angle =80°.

#### Sleep

MRI images were acquired on a 3-T Siemens Trio scanner (Erlangen, Germany). EEG via a cap (modified BrainCapMR, Easycap, Herrsching, Germany) was recorded continuously during fMRI acquisition (1505 volumes of T2-weighted echo planar images, TR/TE = 2080 ms/30 ms, matrix 64×64, voxel size 3×3×2 mm3, distance factor 50%; FOV 192 mm^2^). An optimized polysomnographic setting was employed (chin and tibial EMG, ECG, EOG recorded bipolarly [sampling rate 5 kHz, low pass filter 1 kHz] with 30 EEG channels recorded with FCz as the reference [sampling rate 5 kHz, low pass filter 250 Hz]. Pulse oxymetry and respiration were recorded via sensors from the Trio [sampling rate 50 Hz]) and MR scanner compatible devices (BrainAmp MR+, BrainAmpExG; Brain Products, Gilching, Germany), facilitating sleep scoring during fMRI acquisition.

#### Disorders of consciousness: Paris

MRI images were acquired with two different acquisition protocols. In the first protocol, MRI data of 26 patients and 13 healthy controls were acquired on a 3T General Electric Signa System. T2*-weighted whole brain resting state images were acquired with a gradient-echo EPI sequence using axial orientation (200 volumes, 48 slices, slice thickness: 3 mm, TR/TE: 2400 ms/30 ms, voxel size: 3.4375 × 3.4375 × 3.4375 mm, flip angle: 90°, FOV: 220 mm^2^). An anatomical volume was also acquired using a T1-weighted MPRAGE sequence in the same acquisition session (154 slices, slice thickness: 1.2 mm, TR/TE: 7.112 ms/3.084 ms, voxel size: 1 × 1 × 1 mm, flip angle: 15°).

In the second protocol, MRI data of 51 patients were acquired on a 3T Siemens Skyra System. T2*- weighted whole brain resting state images were recorded with a gradient-echo EPI sequence using axial orientation (180 volumes, 62 slices, slice thickness: 2.5 mm, TR/TE: 2000 ms/30 ms, voxel size: 2 × 2 × 2 mm, flip angle: 90°, FOV: 240 mm^2^, multiband factor: 2). An anatomical volume was acquired in the same session using a T1-weighted MPRAGE sequence (208 slices, slice thickness: 1.2 mm, TR/TE: 1800 ms/2.35 ms, voxel size: 0.85 × 0.85 × 0.85 mm, flip angle: 8°).

#### Disorders of consciousness: Liège

MRI images were acquired on a Siemens 3T Trio scanner (Siemens Inc, Munich, Germany). MRI acquisition included a gradient echo-planar imaging (EPI) sequence (32 transversal slices, 300 volumes, TR/TE = 2000 ms/30 ms, flip angle = 78°, voxel size = 3×3×3 mm, FOV = 192 mm); a structural T1 (120 transversal slices, TR = 2300 ms, voxel size = 1.0×1.0×1.2 mm, flip angle = 9°, FOV = 256 mm).

#### Brain parcellation

We used the Schaefer parcellation with 1000 brain areas, based on estimation from a large dataset (n = 1489) (Schaefer et al., 2018), to extract the time series from each subject. Furthermore, we estimated the Euclidean distances from the Schaefer parcellation in MNI space.

### Resting-state pre-processing

#### Datasets: Meditation, Paris, Liège

The pre-processing of resting-state data was performed using FSL (http://fsl.fmrib.ox.ac.uk/fsl) as described in our previous study (López-González et al., 2021). In brief, resting-state fMRI was computed using MELODIC (Multivariate Exploratory Linear Optimized Decomposition into Independent Components) (Beckmann & Smith, 2004). Steps included discarding the first five volumes, motion correction using MCFLIRT (Jenkinson et al., 2002), Brain Extraction Tool (BET) (Smith, 2002), spatial smoothing with 5 mm FWHM Gaussian Kernel, rigid-body registration, high pass filter cutoff = 100.0 s, and single-session ICA with automatic dimensionality estimation. Then, lesion-driven artifacts (for patients) and noise components were regressed out independently for each subject using FIX (FMRIB’s ICA-based X-noiseifier) (Griffanti et al., 2014). Finally, FSL tools were used to co-register the images and extract the time-series between 1000 cortical brain areas for each subject in MNI space from the Schaefer parcellation (Schaefer et al., 2018).

#### Sleep

The pre-processing of resting-state data was performed using FSL (http://fsl.fmrib.ox.ac.uk/fsl). In brief, steps included discarding the first five volumes, motion correction using MCFLIRT (Jenkinson et al., 2002), BET (Smith, 2002), spatial smoothing with 5 mm FWHM Gaussian Kernel, rigid-body registration, bandpass filtering between 0.01 − 0.1 *Hz*. Finally, FSL tools were used to co-register the images and extract the time-series between 1000 cortical brain areas for each subject in MNI space from the Schaefer parcellation (Schaefer et al., 2018). Previous publications based on this dataset can be consulted for further details (Perl, Pallavicini, et al., 2021).

#### Probabilistic Tractography analysis

We used the Human Connectome Project (HCP) database that contains diffusion spectrum and T2-weighted neuroimaging data from 32 participants as reported in Deco and Kringelbach (Deco & Kringelbach, 2020). A complete description of the acquisition parameters is described in detail on the HCP website (Setsompop et al., 2013). The freely Lead-DBS software package (https://www.lead-dbs.org/) provides the pre-processing described in detail in Horn et al. (Horn et al., 2017). In brief, the data were processed by using a q-sampling imaging algorithm implemented in DSI studio (http://dsi-studio.labsolver.org). A white-matter mask was computed by segmenting the T2-weighted images and co-registering the images to the b0 image of the diffusion data using SPM12. For each HCP participant, 200,000 fibres were sampled within the white-matter mask. Fibres were transformed into MNI space using Lead-DBS Horn and Blankenburg (Horn & Blankenburg, 2016). Finally, we used the standardized methods in Lead-DBS to extract the structural connectomes from the Schaefer 1000 parcellation (Schaefer et al., 2018).

### Model-free framework

#### Kuramoto Local order parameter

The amplitude turbulence, 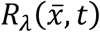, is defined as the modulus of the Kuramoto local order parameter for a given brain area as a function of time:

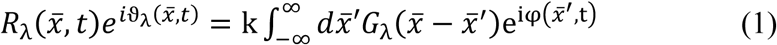

where *G*_*λ*_ is the local weighting kernel 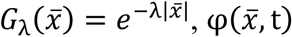 are the phases of the spatiotemporal data and k is the normalisation factor 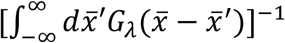. Note that the spatial scaling is defined by *λ*.

Thus, *R*_*λ*_ defines local levels of synchronisation at a given scale, *λ*, as function of space, 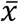, and time, *t*. This measure captures what we here call *brain vortex space, R*_*λ*_, over time, inspired by the rotational vortices found in fluid dynamics, but of course not identical.

#### Amplitude turbulence

The level of amplitude turbulence, *D*, is defined as the standard deviation across time and space of the modulus of local Kuramoto order parameter (R):

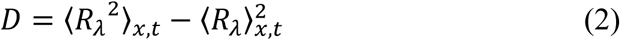

where the brackets ⟨ ⟩_*x,t*_ denotes averages across time and space.

#### Information cascade flow and Information cascade

The information cascade flow indicates how travels the information from a given scale (*λ*) to a lower scale (*λ* − Δ*λ*, where Δ*λ* is a scale step) in consecutive time steps (*t* and *t* + Δ*t*). In this sense, the information cascade flow measures the information transfer across scales computed as the time correlation between the Kuramoto local order parameter in two consecutive scales and times:

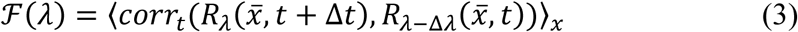

where the brackets ⟨ ⟩_*x,t*_ denotes averages across time and space. Then, the information cascade is obtained by averaging the information cascade flow across scales *λ*, which captures the whole behaviour of the information processing across scales (**Fig. 1A**, middle panel).

#### Information transfer

The spatial information transfer indicates how the information travels across space at a specific scale, *λ*. This measurement is computed as the slope of a linear fitting in the log-log scale of the time correlation between the Kuramoto local order parameter of two brain areas at the same scale as a function of its Euclidean distance (*r*) within the inertial subrange (**Fig. 1A**, right panel).

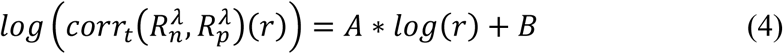

where *A* and *B* are the fitting parameters, and the first one, the negative slope, stands for the spatial information transfer.

#### Node variability of local synchronisation: node-level turbulence

We computed the node variability of the local synchronisation as the standard deviation across time of the local Kuramoto order parameter as follows:

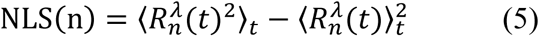

where the brackets ⟨ ⟩_*t*_ represent average values across time points.

Here, we used the discrete version of the node-level Kuramoto order parameter, with modulus *R* and phase *ν*, representing a spatial average of the complex phase factor of the local oscillators weighted by the coupling calculated through:

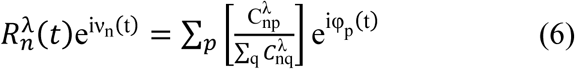

where *ϕ*_*p*_(*t*) are the phases of the spatiotemporal data and 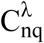 is the local weighting kernel between node *n* and *p*, and *λ* defines the spatial scaling:

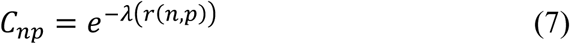

where *r*(*n,q*) is the Euclidean distance between the brain areas *n* and *p* in MNI space.

To compare the node-level turbulence statistics, we collected the 1000 nodes values for all participants in each condition and generated the distributions. Then, we compared across states the distributions using the Kolmogorov-Smirnov distance between them. The Kolmogorov–Smirnov distance quantifies the maximal difference between the cumulative distribution functions of the two samples, where larger values stand for more significant differences between both distributions.

### Model-based framework

#### Whole-Brain Computational Model

We constructed whole-brain dynamical models based on the normal form of a supercritical Hopf bifurcation (also known as Stuart-Landau) (Deco, Kringelbach, et al., 2017). This type of bifurcation can change the qualitative nature of the solutions from a limit cycle that yields self-sustained oscillations towards a stable fixed point in phase space. This whole-brain computational model is characterised by a series of model parameters that rules the global dynamical behaviour. One of them is the multiplicative factor, *G*, representing the global conductivity of the fibres scaling the structural connectivity between brain areas, which is assumed to be equal across the brain (Deco, Kringelbach, et al., 2017; Deco, Tagliazucchi, et al., 2017). The other relevant parameters are the local bifurcation parameter (*a*_*j*_), which rules the dynamical behaviour of each area between noise-induced (*a* < 0), self-sustained oscillations (*a* > 0) or a critical behaviour between both (*a* ∼ 0) (**Fig.1C**). We optimized the model parameters to better fit the empirical functional connectivity as a function of the distance, *r*, within the inertial subrange. The models consisted of 1000 cortical brain areas from the resting-state atlas mentioned above. The underlying anatomical matrix *C*_*np*_ was added to link the brain structure and functional dynamics and was obtained by measuring the exponential distance rule as defined in Equation (7). The local dynamics of each brain area was described by the normal form of a supercritical Hopf bifurcation, which emulates the dynamics for each brain area from noisy to oscillatory dynamics as follows:

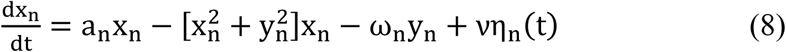

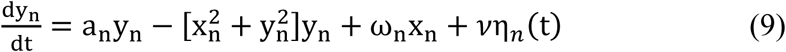

where *η*_*n*_(*t*) is additive Gaussian noise with standard deviation *ν* = 0.01. This normal form has a supercritical bifurcation at *a*_*n*_ = 0, such that for *a*_*n*_ > 0, the system is in a stable limit cycle oscillation with frequency *f*_*n*_ = *ω*_*n*_/2*π*, whereas for *a*_*n*_ < 0, the local dynamics are in a stable point (i.e., noisy state). The frequency *ω*_*n*_ of each brain area was estimated from the empirical fMRI data as the peak of the power spectrum.

Finally, the whole-brain dynamics was defined by the following set of coupled equations:

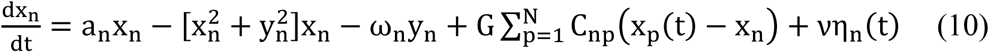

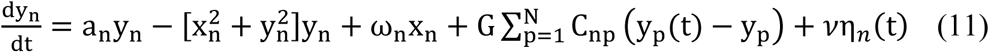

Where the global coupling factor G, scaled equally for each brain area, represents the input received in region *n* from every other region *p*.

#### Functional Connectivity Fitting

Kolmogorov’s structure-function of a variable *u* was applied to the BOLD signal of the data. This measure is based on the functional correlations between each pair of brain areas with equal Euclidean distance and was defined as:

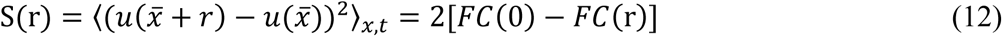

where *FC*(*r*) is the spatial correlations of two points separated by a Euclidean distance r, which is given by:

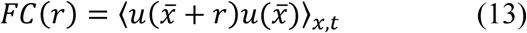

where the symbol ⟨ ⟩_*x,t*_ refers to the average across the spatial location *x* of the brain areas and time. Thus, the structure functions characterise the evolution of the functional connectivity (FC) as a function of the Euclidean distance between equally distant nodes, which is different from the usual definition of FC that does include distance. We then compute the fitting as the Euclidean distance between simulated and empirical FC(r) within the inertial range as defined in Deco et al. (Deco & Kringelbach, 2020).

#### Susceptibility

The susceptibility measure of the whole-brain model was defined as the brain’s sensitivity to react to external stimulations. The Hopf model was perturbed for each G by randomly changing the local bifurcation parameter, *a*_*n*_, in the range [-0.02: 0]. The sensitivity of the perturbations on the spatiotemporal dynamics was calculated by measuring the modulus of the local Kuramoto order parameter as:

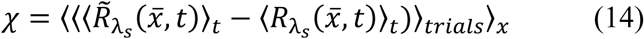

Where 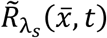 corresponds to the perturbed case, the 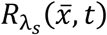 to the unperturbed case, and ⟨ ⟩_*t*_,

⟨ ⟩_*trials*_ and ⟨ ⟩_*x*_ to the average across time, trials, and space, respectively.

#### Information encoding capability

The information encoding capability captures how the external stimulations are encoded in whole-brain dynamics. The information capability, I, was defined as the standard deviation across trials of the difference between the perturbed 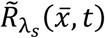 and unperturbed 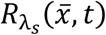 mean of the modulus of the local Kuramoto order parameter across time *t*, averaged across all brain areas *n* as:

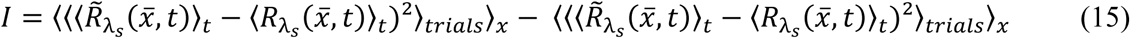

where the brackets ⟨ ⟩_*t*_, ⟨ ⟩_*trials*_ and ⟨ ⟩_*x*_ denote the averages defined as above.

### Statistical Analyses

We applied the Wilcoxon rank-sum method to test the differences between conditions in turbulence, information capacity, information transfer, and perturbative measures. For the node-level analysis, we applied the Kolmogorov–Smirnov test to compare between conditions. Additionally, we applied the False Discovery Rate (FDR) at the 0.05 level of significance to correct multiple comparisons (Hochberg & Benjamini, 1990).

## Acknowledgements

A.E and Y.S.P. are supported by the HBP SGA3 Human Brain Project Specific Grant Agreement 3 (grant agreement no. 945539), funded by the EU H2020 FET Flagship programme. Y.S.P is supported by European Union’s Horizon 2020 research and innovation programme under the Marie Sklodowska-Curie grant 896354. G.D. is supported Spanish national research project (ref. PID2019-105772GB-I00 MCIU AEI) funded by the Spanish Ministry of Science, Innovation and Universities (MCIU), State Research Agency (AEI); HBP SGA3 Human Brain Project Specific Grant Agreement 3 (grant agreement no. 945539), funded by the EU H2020 FET Flagship programme; SGR Research Support Group support (ref. 2017 SGR 1545), funded by the Catalan Agency for Management of University and Research Grants (AGAUR); Neurotwin Digital twins for model-driven non-invasive electrical brain stimulation (grant agreement ID: 101017716) funded by the EU H2020 FET Proactive programme; euSNN European School of Network Neuroscience (grant agreement ID: 860563) funded by the EU H2020 MSCA-ITN Innovative Training Networks; CECH The Emerging Human Brain Cluster (Id. 001-P-001682) within the framework of the European Research Development Fund Operational Program of Catalonia 2014-2020; Brain-Connects: Brain Connectivity during Stroke Recovery and Rehabilitation (id. 201725.33) funded by the Fundacio La Marato TV3; Corticity, FLAG–ERA JTC 2017 (ref. PCI2018-092891) funded by the Spanish Ministry of Science, Innovation and Universities (MCIU), State Research Agency (AEI). MLK is supported by the Center for Music in the Brain, funded by the Danish National Research Foundation (DNRF117), and Centre for Eudaimonia and Human Flourishing at Linacre College funded by the Pettit and Carlsberg Foundations. The study was supported by the University and University Hospital of Liège, the Belgian National Funds for Scientific Research (FRS-FNRS), the European Space Agency (ESA) and the Belgian Federal Science Policy Office (BELSPO) in the framework of the PRODEX Programme, the BIAL Foundation, the Mind Science Foundation, the fund Generet of the King Baudouin Foundation, the Mind-Care foundation and AstraZeneca Foundation. RP is research fellow, OG is research associate, and SL is research director at FRS-FNRS. The authors thank all the patients and participants, the whole staff from the Radiodiagnostic and Nuclear departments of the University Hospital of Liège.

## Author contributions

Conceptualization: AE, YS, MLK, GD. Methodology: AE, YS, MLK, GD. Data analysis: AE, YS. Visualization: AE, YS, MLK, GD. Supervision: MLK, GD. Data preprocessing: AE, CP, ALG, MLK. Data curation: EC, BT, NP, RP, JA, OG, SL, LN, JDS, HL, ET, MLK. Writing—original draft: AE, YS, MLK, GD. Writing—review & editing: CU, EC, ALG, JDS, ET, OG, LN.

## Figures

**Fig. S1:**
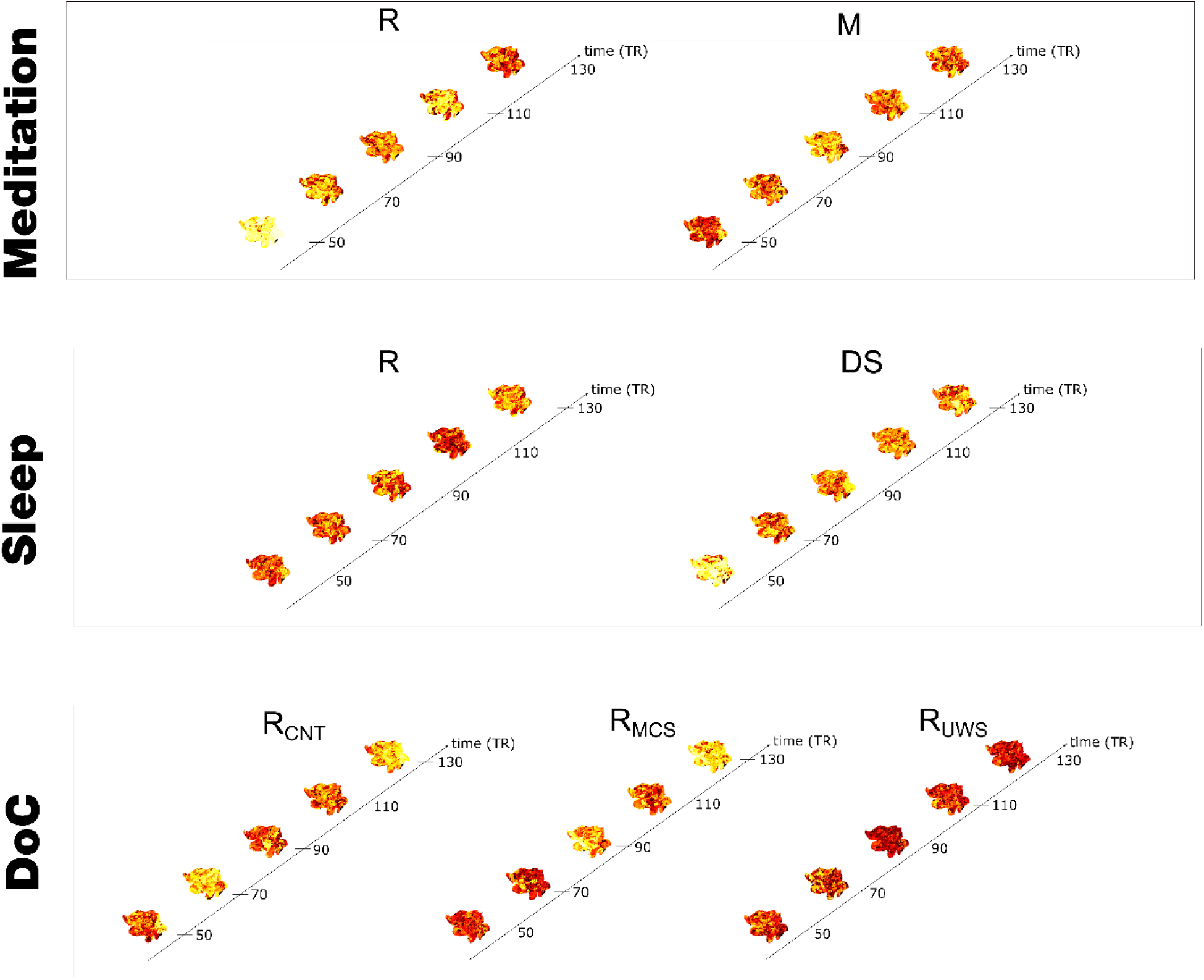
Spatiotemporal evolution of turbulent in empirical data. Continuous snapshots for segments separated in time (20 TRs) rendered on a flatmap of the hemisphere for each condition within each dataset.

